# Goal-Dependent Hippocampal Representations Facilitate Self-Control

**DOI:** 10.1101/2021.08.26.457750

**Authors:** Micah G. Edelson, Todd A. Hare

**Affiliations:** Zurich Center for Neuroeconomics, Department of Economics, University of Zurich, 8006 Zürich, Switzerland

## Abstract

Mnemonic influences on decision-making processes are important for linking past experiences and simulations of the future with current goals. The ways in which mnemonic information is represented may be especially critical in situations where one needs to overcome past rewarding experiences and exert self-control. We propose that self-control success or failure may depend on how information is retrieved from memory and how effectively this memory retrieval process can be modified to achieve a specific goal. We test our hypothesis using representational similarity analyses of human neuroimaging data during a dietary self-control task in which individuals must overcome taste temptations to choose healthy foods. We find that self-control is indeed associated with the way individuals represent taste information in the brain and how taste representations adapt to align with different goals. These results provide new insights into the processes leading to self-control success and indicate the need to update the classical view of self-control to continue to advance our understanding of its behavioral and neural underpinnings.

## Introduction

The capability to adjust our thoughts and behavior to overcome temptations and pursue a higher order goal is central to human success and well-being. Despite its importance and a long history of inquiry, the underlying mechanisms leading to successful self-control are still only partially understood. Longstanding theories of the neurobiology of self-control have primarily focused on regulatory functions supported by the prefrontal cortex and antagonistic interactions between the prefrontal cortex and subcortical brain regions associated with reward such as the amygdalae and striatum (1–8). However, self-control challenges often require individuals to assess past experiences (rewarding or aversive) and use this information to simulate the effects of current actions. Therefore, another potentially important, yet thus far overlooked, component in self-control success is mnemonic processing.

We hypothesized that one key to self-control success is to adapt the sources of past information that inform the choice so that they align with the current goal. Such a role for memory in self-control would be consistent with the growing body of work on memory sampling theories (9–11) and empirical data from human and other animal models on the relationship between memory and decision making (12–18). To test this hypothesis at the neurobiological level, we focused on the hippocampus because it is theorized to be a prominent source of information and simulations of future outcomes for goal-directed decisions (13, 19) and is the region most closely associated with episodic memory retrieval and sampling in past research (9, 10, 20–23). We predicted that representations of stimuli and their component attributes in the hippocampus should adjust to current goals, and that self-control success will depend on the extent to which brain representations of attribute-level information align with the current goal.

We tested our hypothesis using human neuroimaging and behavioral data during acquired during food evaluation and consumption choice tasks. We found that multidimensional hippocampal representations of food items change to align with task conditions and goals. Specifically, greater adaptations of hippocampal representations of the palatability of food items to a self-control goal were associated with healthier consumption choices.

## Results

The participants in this study were self-reported dieters who evaluated the same fifty food items in the context of three separate goals while undergoing functional magnetic resonance imaging (Figure 1). To examine the multi-faceted goal directed representational patterns of episodic memory traces we used representational similarity analyses (RSA), a technique well suited to capture such patterns (24–26, 20). RSA quantifies the patterns of neural activity in a given region (i.e., the relative activation levels of each voxel in the region) and can then test if this pattern changes under different goals or how well it aligns with a theoretical model. For example, this approach allows us to test whether brain activity patterns distinguish between subjectively tasty and non-tasty food items, and whether the current goal can influence this distinction.

**Figure 1.**
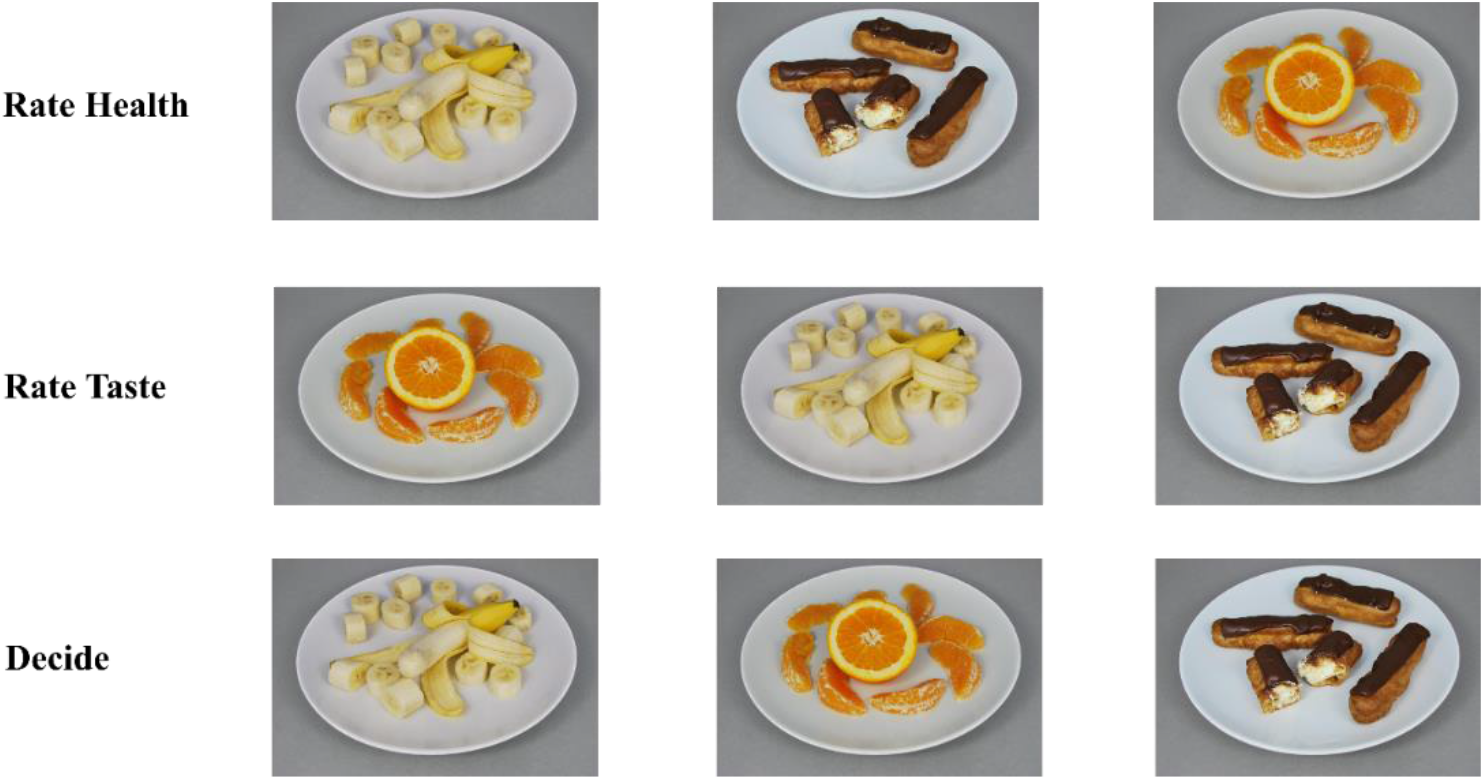
Task structure. Participants evaluated the same 50 food items under three distinct goals. They rated the palatability of each food item independently of its healthiness (goal = evaluate taste). They also rated the healthiness of each food item independently of its palatability (goal = evaluate healthiness) (counterbalanced). Lastly, they chose whether they would eat the food item currently shown on the screen or a fixed default food item (goal = choose the subjectively best option to consume). Ratings and choice were done on a 5-point scale and one randomly selected item was actually consumed by participants in the end of the experiment.

We first tested if the different task-induced goals influenced the representations of the same food items within the hippocampus. To do so we tested if the neural activity patterns showed stronger support for a model that assumes that items are represented similarly across goals vs. a model that assumes that hippocampal representations shift across goals (**Figure 2**; Goal vs. Trial based model comparison, Tau difference=0.035; *p*<0.0000001; via Kendall rank correlation test which is the standard RSA measure for testing the ordinal association between a given model and brain activity patterns). These results in human participants are consistent with previous findings in rodents showing that the representation in several regions, including the hippocampus change across motivational states and task contingencies (27–29). These previous findings demonstrate an overall effect of goals on hippocampal representations (which could be driven by many things, e.g., attention, motivation), but do not test whether the hippocampal representations are associated with self-control.

**Figure 2.**
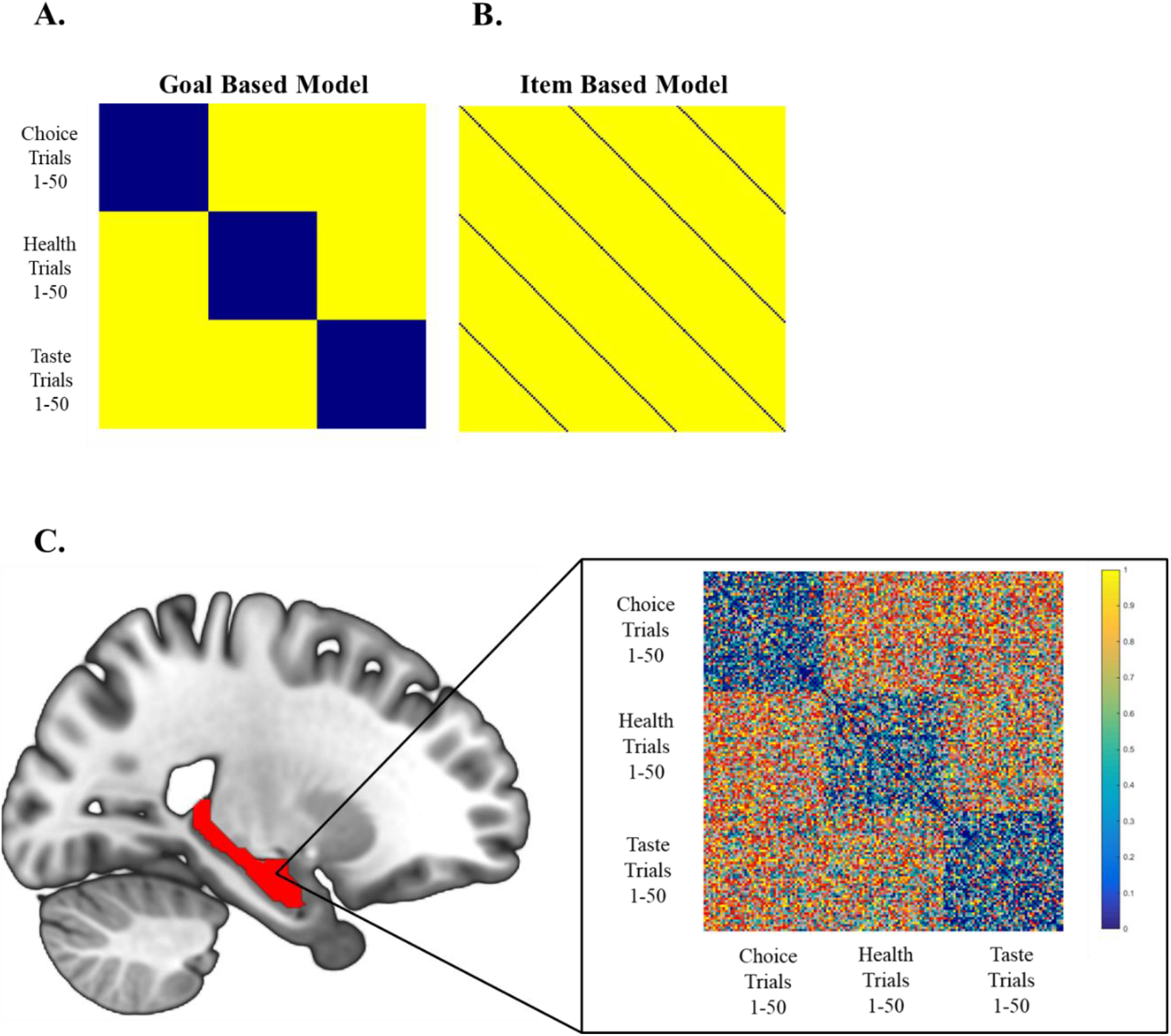
Hippocampal representations of food items are goal dependent. **A-B.** The two matrices on top show theoretical models describing either a high similarity in representations across the 50 foods under the same goals, or alternatively across repeated presentations of the same items regardless of the behavioral goal. All matrices depict the dissimilarity (1 – Spearman correlation, with values scaled to range from 0 - 1) between trials, and thus blue indicates greater similarity **A.** In the goal-dependent model, the 50 row by 50 column blocks in blue along the diagonal from upper left to bottom right indicate the hypothesized high degree of representational similarity across the 50 different foods under the same goal. In contrast, the off-diagonal blocks in yellow indicate the predicted high degree of dissimilarity between the same food items under different task goals. **B.** In the item-based model, the diagonal lines of blue squares show its prediction of high similarity between presentations of the same food item regardless of the behavioral goal. **C.** The matrix depicts the group average dissimilarity (i.e., 1-Spearman correlation) in brain activity across all voxels in the hippocampal region of interest (ROI, shown in red on the brain image to the left). The 50 × 50 blue square patterns with low dissimilarity along the diagonal and generally high dissimilarity values in other locations indicate that the task goal strongly influenced the nature of food items representations in the hippocampus (Goal vs. Trial based model comparison, Tau alpha difference=0.035; p <0.0000001).

### Hippocampal representations of taste are flexible and goal-dependent

Given that hippocampal representations for the same items are goal dependent (**Figure 2**), we hypothesized that hippocampal activity would clearly distinguish between high and low palatability during taste ratings, but that this difference would be less distinct when the goal is not dependent on taste (e.g., when making health ratings) (**Figure 3 A-B**). Indeed we find that the dissimilarity in hippocampal representations of more, relative to less, palatable items is reduced in the health rating sessions compared to both the taste rating and decision sessions (**Figure 3C**; Cohen’s d = 0.69, posterior probability of difference > 0 = 0.999; Cohen’s d = 0.50, posterior probability of difference > 0 = 0.992, respectively).

**Figure 3.**
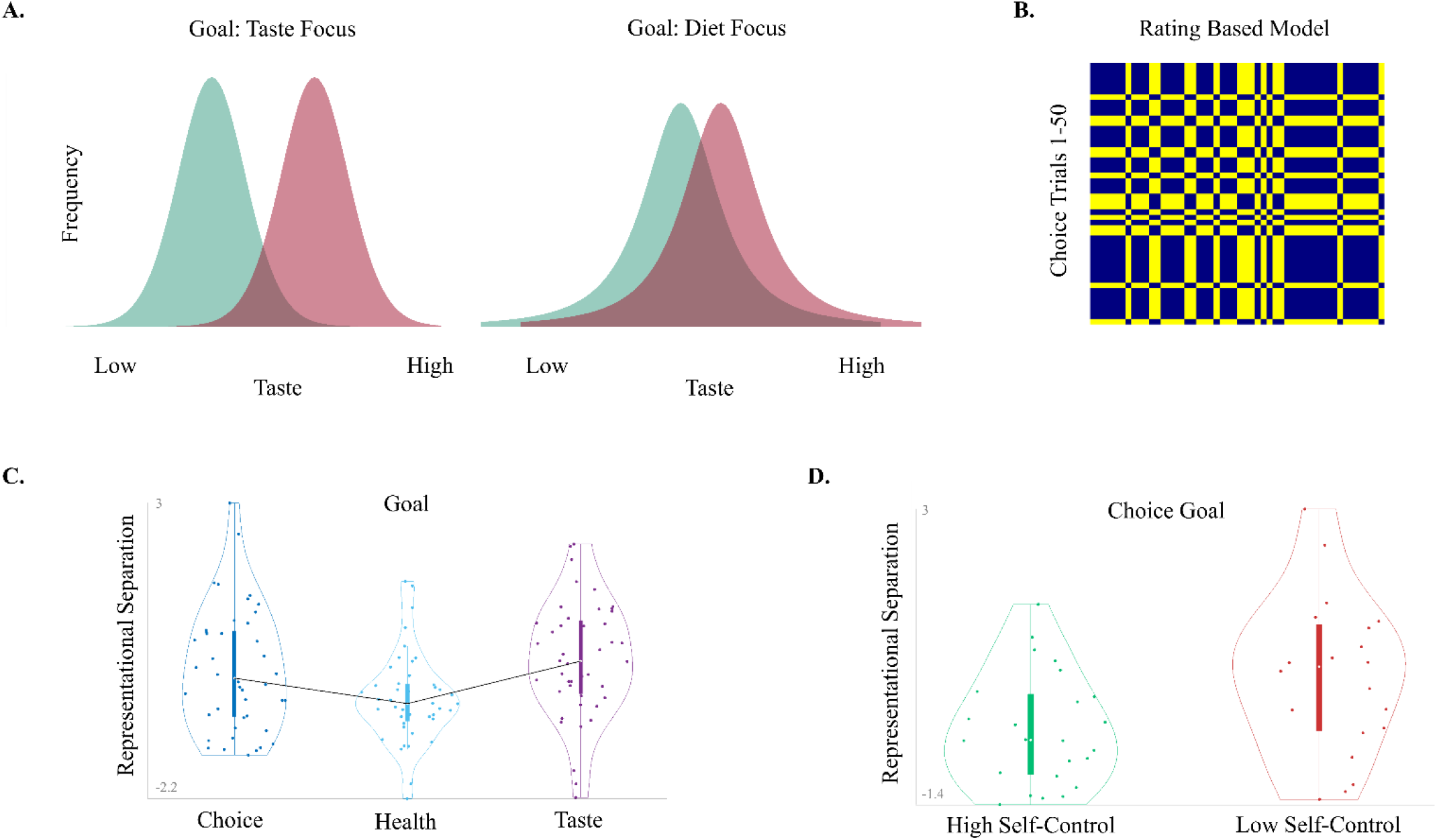
Hippocampal representations of food palatability are related to dietary self-control. **A.** Illustrative example for a goal dependent change in the distinctness of high vs. low taste items. The colored distributions represent theoretically sampled subsets of past experiences with two example food items, one with a high mean value for taste (Red) and one with a low mean value for taste (Green). When the goal is to focus on tastiness (left panel), a sample of past consumption experiences and/or predictions is drawn such that the tastier item is clearly distinct from the less tasty item. Under a goal to exert self-control (right panel) a larger overlap between the tastier and less tasty items may occur do to 1) biased sampling of past events that favors less positive samples of the highly tasty food and more positive samples of less tasty food, and/or 2) increased noise in the sampling process because less attention or other resources are devoted to taste attributes leading to wider distributions. A change from the left to right pattern should facilitate choosing the less tasty, but healthier items more often under the self-control goal. **B.** An example of a single participant’s theoretical model separating high vs. low ratings in the choice task (i.e., pairs of items that were both rated similarly, either high or low, will be depicted in blue). Note that because each participant has different preferences for different food items, each participant’s model is unique. **C.** Hippocampal taste representations depend on the behavioral goal. Hippocampal representations distinguish tastiness to a larger degree when the goal includes the need to evaluate tastiness (i.e. during the taste and choice phases) than when the goal is to ignore taste and concentrate only on healthiness (Taste vs. Heath phase mean standardized difference (Cohen d) = 0.69, posterior probability of an effect larger than 0 = 99.9%; Choice vs. Health phase: mean standardized difference (Cohen d) =0.5, posterior probability of an effect larger than 0 =99.9%). **D.** High self-controllers do indeed have less separation between their representations of palatable and unpalatable foods, relative to low self-controllers when making consumption decisions (Cohen’s d = 0.8, posterior probability low > high self-controllers = 0.99).

### Hippocampal taste representations at the time of choice are associated with successful self-control

We next tested if hippocampal representations contain information about the healthiness and taste of each food item and how these representations relate to actual self-control choices. Indeed, when dieters need to choose to eat food items, there are distinct multi-voxel patterns of hippocampal activity for healthy versus unhealthy foods (peak searchlight center = −18 −33 9; pseudo t = 7.3, p < 0.0002, FWE corrected) as well as more palatable versus less palatable foods (peak searchlight center = −36 −15 −12; pseudo t = 6.1, p < 0.0002, FWE corrected). However, only the degree of separation related to the tastiness of the food items was strongly associated with higher self-control success (see supplementary results for details on healthiness representations). We define self-control success as choosing a healthier, but less tasty item over a tastier, but less healthy option, and classified individuals that successfully employed self-control more than 50% of the time as high self-controllers and those with less than 50% success as low self-controllers. When making consumption decisions, high self-controllers had less separation between their representations of palatable and unpalatable foods than low self-controllers (Cohen’s d = 0.80, posterior probability low > high self-controllers = 0.989, **Figure 3D**). Thus, in line with memory sampling theories (9–11), our results indicate that one key to self-control may be selecting the types of mnemonic information that are prioritized and sampled from, when making a choice (not necessarily consciously).

Note that although both the choice and taste rating tasks require one to evaluate taste information, self-control only needs to be exercised during the choice task. For successful self-control it may be beneficial to sample from memory the subset of events where very tasty items (e.g., chocolate cake) were less tasty than average, and the subset of events were less tasty items (e.g., cabbage) were unusually appetizing. In other words, under a self-control goal the difference between representations of highly tasty and less tasty items will decrease, or become less distinct, when compared to the same contrast when the goal is to evaluate the items’ tastiness (**Figure 3A**).

We constructed a simple classifier to test this hypothesis. The classifier uses the degree of separation in the hippocampal taste representations in the decision versus taste rating tasks to predict the self-control level for each individual using a leave-one-out approach (see Methods). The change in hippocampal taste representations significantly predicted whether an individual would have high versus low self-control success (accuracy=0.68, probability results are greater than chance = 0.996).

Finally, although our primary focus in this work was on examining the relationship between hippocampal representations and self-control, we ran an additional exploratory whole brain searchlight analysis testing for brain regions in which changes in the representational similarity of palatable and unpalatable foods between the taste-rating and choice phases, differed between individuals with high and low self-control success (FWE<0.01). We found a single region in the striatum (Left Caudate, MNI −18 24 0, pseudo t= 5.28 FWE=0.003, k=17) that showed a greater distinction between palatable and unpalatable foods during the decision session in low relative to high self-controllers. Interestingly, the striatum is implicated in associative memory processes as well as in signaling reward affect, anticipation, craving, motivation, and outcomes (29–33). Changes in any of these processes between the taste-rating and choice phases could plausibly lead to more successful dietary self-control.

## Discussion

The way the brain retrieves choice-relevant information from past experiences may play a key role in the ability to modify or regulate the decision process. The hippocampus is thought to be a key source of evidence used to construct subjective values and take goal-directed decisions (12–18, 34). Here, we show that goal-dependent changes in hippocampal representations are associated with the ability to exert self-control during dietary choices.

Memory sampling processes may be an important component of self-control. Theories of memory retrieval propose that it is an estimation based on repeated sampling of noisy internal representations. These theories can explain mnemonic phenomenon such as context effects as well as regularities in choice and decision time (9, 10, 14) while accounting for the way neural populations encode information (35). There are multiple plausible mechanisms that might lead to goal-dependent hippocampal representations of palatability. The current goal may determine the accessibility of specific attributes such that when taste attributes are irrelevant or inconsistent with the goal they are sampled less often or less precisely. The goal may also influence the types of past experiences that are retrieved or imagined. For example, if the goal is to eat healthfully, then instances when chocolate cake was dried out and unpleasant or of an unexpectedly tasty broccoli dish may be sampled more often. This shift in the type of information sampled could lead to less distinct representations (**Figure 3A**). These and other attention-related processes may combine to alter the way individuals recall and/or imagine the taste of foods in different goal contexts, thereby facilitating goal-consistent choices.

Our results are consistent with choice theories that propose that preferences are not stored in an immutable format, but rather constructed at the time of choice and can be influenced by cognitive and contextual factors (26, 36–38). Cognitive reappraisal, or the modification of the way one thinks about a stimulus in order to change its emotional or other subjective evaluations (33, 39–41), is a form of goal-dependent evaluation of previous and current experiences. Concurrent as well as previously trained reappraisal of food and drug stimuli reduce craving scores, willingness to pay, and/or consumption of cigarettes and food (1, 33, 42). Furthermore, greater activity in the hippocampal complex during emotion reappraisal correlates with better dietary self-control (43). In regulation of craving training (ROC-T) (42) participants view pictures of food items and positively or negatively reappraise their values. Undergoing ROC-T might alter the distribution of experiences with the food items and/or change the accessibility or ease of retrieval for positive versus negative aspects of the foods. Previous neuroimaging studies of emotion and cigarette reappraisal have focused on interactions between prefrontal cortex and amygdala or ventral striatum during active reappraisal. However, our results suggest that future studies should test if ROC-T leads to changes in hippocampal multivariate representations of tempting attributes (e.g., food taste or cigarette pleasure) even when people are no longer explicitly reappraising the stimuli, whether those representations mediate increases in healthful behavior, and how long the altered representations are maintained after training.

Previous work in healthy and clinical populations has demonstrated that successful self-control is associated with prefrontal cortex activation and its connectivity with subcortical areas such as amygdala, and striatum (1–8). Our current findings indicate the need to expand this classical view of control networks to incorporate hippocampal representations as well. It remains to be seen whether the hippocampal complex interacts with the same prefrontal regions previously linked to self-control or if instead it operates as part of a different network.

Self-control is known to be an important factor in both physical and mental health as well as educational and social achievements (6, 44, 45). Our results provide new insights into the behavioral and neurobiological processes underling self-control and indicate the need to incorporate mechanistic frameworks of memory representations to further advance our understanding of this critical capacity. These insights could lead to new ways to approach treatment of self-control-related disorders or maximize self-control skills in healthy individuals.

## Methods

### Participants

We used a dataset originally reported in Hare et., al. (2). In total, fifty-two participants completed the experiment. All participants were right-handed, healthy, had normal or corrected-to-normal vision, had no history of illnesses, and were not taking medications that interfere with the performance of fMRI. Participants had no history of eating disorders or food allergies to any of the items used in the experiment. Participants were told that the goal of the experiment was to study food preferences among dieters and gave written consent before participating. The initial study recruited two types of participants: 1) individuals who self-reported being on a diet to lose or maintain weight, and 2) individuals who self-reported no current monitoring of their diet. All participants reported that they enjoyed eating sweets, chocolate, and other “junk food” even though they might be restricting them from their current diet. The review board of the California Institute of Technology approved the study. Three participants were excluded due to multiple large head movements exceeding 3 mm. Five participants were also excluded because the RSA toolbox (25) quality assurance measures indicated that the default correlation metric was not appropriate for their data due to ‘small relative standard deviations’. As a conservative measure, we opted to simply exclude these participants from the analyses rather than exploring different distance matrix options. Note that in the original study some participants were excluded from the analyses based on a set of *a priori* criteria regarding choice behavior. However, in the current paper, we analyzed the data without imposing any behavior-based exclusion criteria, leaving us with 44 participants in the analyses reported here.

### Stimuli and task

We briefly summarize the relevant details of the task here; additional details can be found in Hare et., al (2). Participants rated and made decisions on 50 different food items including junk foods (e.g., chips or candy bars) as well as healthy snacks (e.g., apples or broccoli). For visual illustration purposes in Figure1A, high resolution images were taken from an online food image database (46). Participants were instructed not to eat for three hours before the experiment, which is known to increase the value that is placed on food. The task had three parts, all of them done in the scanner. Participants first rated all 50 food items for both their taste and healthiness in two separate blocks (a taste-rating block and a health-rating block). The order of the rating blocks was counterbalanced across participants and the food items were presented in random order. Ratings were made using a five-point scale that was shown on the screen below each item. The health rating scale was: Very Unhealthy, Unhealthy, Neutral, Healthy, Very Healthy. The taste rating scale was: Very Bad, Bad, Neutral, Good, Very Good. The mapping of rating labels to button orders was counterbalanced across participants. Before the taste-rating block participants were instructed to “rate the taste of each food item without regard for its healthiness”. Before the health-rating block they were instructed to “rate the healthiness of each food item without regard for its taste”. Participants had a maximum of 4 seconds to enter their rating and the trial terminated as soon as they did so. Trials were separated by a random ITI with duration distributed uniformly between 4 and 15 seconds. Following the two rating blocks, one item that was rated as neutral on both health and taste was selected as the reference food for that participant used in the choice task (see below). Examples of such reference items included wheat crackers, jello, raisins, granola bars, and yogurt.

The final session of the experiment was a decision phase. At the beginning of this phase participants were shown a picture of the reference item and told that on each trial they would have to choose between eating the food item shown in that trial and the reference food. Participants had a maximum of 4 seconds to enter their decision and the trial terminated as soon as they did so. Trials were separated by a random ITI with duration distributed uniformly between 4 and 15 seconds. Each food item was shown once for a total of 50 decision trials. Participants were required to eat the food that they chose in a randomly selected trial at the end of the experiment. Although this was a binary decision task, participants were asked to express the strength of their preferences using a five-point scale: Strong No (=choose reference), No (=choose reference item), Neutral, Yes (=choose shown item), Strong Yes (=choose shown item). In the trials in which the participants choose ‘Neutral’ a coin was flipped to determine the decision.

### Self-control success definition

For each participant we calculated the percentage of conflict trials (i.e., trials in which one choice option was healthier but the other was tastier), where the healthier option was chosen. In order to maintain consistency with previous work(2), a neutral decision response was not considered to be a failed self-control event.

### fMRI data acquisition

Functional imaging was conducted using a 3.0 Tesla Siemens Trio MRI scanner to acquire gradient echo T2*-weighted echoplanar (EPI) images with BOLD contrast. Images were acquired in an oblique orientation of 30° to the anterior commissure–posterior commissure line. Each volume of images had 44 axial slices. A total of 777 volumes were collected over three sessions in an interleaved-ascending manner. The imaging parameters were as follows: echo time, 30 ms; field of view, 192 mm; in-plane resolution and slice thickness, 3 mm; repetition time, 2.75 s.

### Image analysis

Statistical Parametric Mapping (SPM 12; Wellcome Trust Centre for Neuroimaging, London, UK, http://www.fil.ion.ucl.ac.uk/spm) standard pipeline was used to pre-process the MRI data. Specifically, images were realigned, unwarped and slice-time corrected (to the middle slice acquisition time). T1-weighted structural images were co-registered with the mean functional image and normalized to the standard T1 template based on the Montreal Neurological Institute (MNI) reference brain, using the segment procedure provided by SPM 12. The functional images were then normalized to a standard EPI template using the same transformation.

For each participant, a time series was created indicating the temporal position of each trial convolved with the canonical hemodynamic response using a random-effects general linear model (GLM). Following the procedure for representational similarity analysis (RSA, outlined in Nili et., al. (25)), each event (i.e. each of the 50 food items) was treated as a separate regressor in a model that included regressors for all 150 trials (50 healthiness ratings, 50 taste ratings, and 50 choices). The six movement regressors generated by SPM were also included as covariates of no-interest in the GLM models. All single participant maps were used as the input to a group-level non-parametric analysis using Statistic non-Parametric Mapping (SnPM 13 toolbox, http://warwick.ac.uk/snpm).

### Region of interest (ROI) definition

The hippocampal ROI was defined anatomically using the automated anatomical labelling atlas (47). We used the left hippocampus because the original findings from Hare et al (2) suggested that activity in a left lateralized circuitry was related to performance in this self-control task.

### Representation similarly analysis (RSA)

First, single-participant searchlight-based dissimilarity maps were computed and used as an input to a non-parametric random-effects group-level analyses implemented in SnPM. To create the single-participant maps we followed the pipeline detailed in the RSA toolbox (25), using the default 15 mm spherical searchlights, with the following exception. When comparing brain activity to a model reflecting the subjective health and taste ratings for each participant, we used a participant-specific rather than a group average model. This was done because the subjective nature of food preferences meant that each participant rated the foods differently and averaging over these values would not be appropriate in this case. The ROI results from the hippocampus are based on average of all spheres centered in voxels in the anatomically defined ROI. To test whether representations in the hippocampus ROI corresponded to a theoretical model separating high (positive) and low (negative and neutral) taste/health ratings, the hippocampal representational dissimilarity matrix (RDM) was compared to a model RDM contrasting non-negative vs. negative ratings (25, 48, 49). To compare goal dependent to trial level similarity, the hippocampal RDM was compared to a theoretical model separating goals or trials respectively (see **Figure 1A**). The resulting tau alpha values quantifying the correspondence between the two model RDMs and the neural RDM were compared directly using the ROI approach implemented in the RSA toolbox (25). Kendall tau-alpha is the statistic from the non-parametric Kendall rank correlation test which is the standard RSA measure for testing the ordinal association or covariance between a given model and a brain activity pattern.

### Bayesian analysis of dissimilarity measures by group and condition

Posterior inferences about the differences between groups (high versus low self-controllers) and conditions (rate taste, rate healthiness, decide) were drawn from 100,000 samples estimated using the default robust priors in the BEST package (50, 51) in R (52) based on an Markov-Chain Monte Carlo (MCMC) process implemented in JAGS (53).

### Leave-one out cross-validation classifier

To quantify if the degree to which individuals change their neural representations when transitioning between goals predicts self-control successes, we used a leave-one out cross-validation approach (54). This procedure first calculates the mean shift in tastiness representations between the taste rating and choice tasks using the data from n-1 participants (training group). Subsequently, the shift in representations for the independent participant that was not included in the n-1 analysis is compared to this value and the participant is assigned to one of two categories. If the left-out individual’s shift in representation is larger than the average training group value, she is predicted to be a high self-controller, and vice versa if the change is smaller than the average group value. An n-fold replication of this procedure produces n predicted (i.e., one for each participant) values that can be compared to the real values of each participant (i.e. are they indeed high or low self-controllers according to their behaviour) to determine average accuracy. This value is then compared to a null distribution created by running the same analysis with shuffled labels 10,000 times to test if it is significantly different from chance.

## Supplementary results

Multivariate patterns in the hippocampus distinguished between healthy and unhealthy items as well as palatable and unpalatable items. In fact, the dissimilarity in representations between high and low health and high and low taste were positively correlated across participants (spearman’s rho= 0.794). However, a Bayesian logistic regression of both taste and health representations against self-control success indicated that larger differences between palatable and unpalatable items at the time of choice were associated with low self-control (standardized coefficient = −0.855, posterior probability < 0 = 0.942), but, when accounting for taste representations, differences between healthy and unhealthy items were unrelated to self-control (standardized coefficient = −0.022, posterior probability < 0 = 0.524).

